# Analysis of the ubiquitin-modified proteome identifies novel host factors in Kaposi’s sarcoma herpesvirus lytic reactivation

**DOI:** 10.1101/2022.08.14.503934

**Authors:** Amerria Causey, Mathew Constantine, Jessica Oswald, Anna Dellomo, Bronwyn Masters, Esosa Omorogbe, Arie Admon, Alfredo Garzino-Demo, Elana Ehrlich

**Affiliations:** Biological Sciences, Towson University, Towson, Maryland, United States; Dept. of Molecular Medicine, University of Padova, Padova, Italy, Dept of Microbial Pathogenesis, University of Maryland School of Medicine, and Dept of Microbiology and Immunology, University of Maryland School of Medicine, Baltimore, MD, USA; Biology, Technion - Israel Institute of Technology, Haifa, Israel

## Abstract

Kaposi’s Sarcoma Herpesvirus (KSHV) is the causative agent of Kaposi’s Sarcoma (KS) and is associated with primary effusion lymphoma (PEL), multicentric Castleman’s disease (MCD) and two inflammatory diseases. KSHV-associated cancers are primarily associated with genes expressed during latency, while other pathologies are associated with lytic gene expression. The major lytic switch of the virus, Replication and Transcription Activator (RTA), interacts with cellular machinery to co-opt the host ubiquitin proteasome system to evade the immune response as well as activate the program of lytic replication. Through SILAC labeling, ubiquitin remnant enrichment and mass spectrometry, we have analyzed the RTA dependent ubiquitin-modified proteome. We identified RTA dependent changes in the populations of polyubiquitin chains, as well as changes in ubiquitinated proteins in both cells expressing RTA and naturally infected cells following lytic reactivation. We observed an enrichment of proteins that are also reported to be SUMOylated, suggesting that RTA, a SUMO targeting ubiquitin ligase, may function to alleviate a SUMO dependent block to lytic reactivation. RTA targeted substrates directly through a ubiquitin ligase domain dependent mechanism as well as indirectly through cellular ubiquitin ligase RAUL. Our ubiquitome analysis revealed an RTA dependent mechanism of immune evasion. We provide evidence of inhibition of TAP dependent peptide transport, resulting in decreased HLA complex stability. The results of this analysis increase our understanding of mechanisms governing the latent to lytic transition in addition to the identification of a novel RTA dependent mechanism of immune evasion.

**Importance:** Kaposi’s sarcoma herpesvirus (KSHV), an AIDS associated pathogen, is associated with multiple cancers and inflammatory syndromes. This virus has a latent and lytic lifecycle, each associated with pathogenesis and oncogenesis. Here we identify proteins that display differential abundance in different phases of the lifecycle. We provide evidence supporting a new model of viral immune evasion. These findings increase our understanding of how the virus manipulates the host cell and provides new targets for intervention.

## Introduction

Kaposi’s Sarcoma Herpesvirus (KSHV), also known as HHV-8, is the causative agent of Kaposi’s Sarcoma (KS) and is associated with primary effusion lymphoma (PEL) and multicentric Castleman’s disease (MCD)(1, 2). KSHV is also linked to the inflammatory diseases, KSHV inflammatory cytokine syndrome and immune reconstitution syndrome-KS (3–5). KSHV is classified as a Group 1 carcinogen by the International Agency for Research on Cancer and the National Toxicology Program 14^th^ Report on Carcinogens (6).

KSHV, a gamma herpesvirus, has a two-phase lifecycle consisting of latency and lytic replication. During latency, few genes are expressed, and genome replication occurs with the purpose of episome distribution to daughter cells during cell division. Lytic replication is initiated via expression of the viral Replication and Transcription Activator (RTA) protein. This results in rapid expression of lytic genes, replication of viral DNA, assembly of virus particles and egress from the cell (7).

KSHV is primarily found in cells as a latent infection. Initiation of KSHV-associated cancers is, in part, associated with the genes expressed during latency, which maintain an environment that allows for the survival of cells harboring latent virus. To maintain a latent infection, several checkpoints and mechanisms of regulation are bypassed to progress through the cell cycle without triggering apoptosis, autophagy, or an antiviral response. Dysregulation of such critical processes as cell cycle control, apoptosis and immune response-related signaling pathways are believed to be associated with development of an oncogenic environment. In addition to virus propagation and proliferation, lytic gene expression is also thought to be required for tumorigenesis in KS and MCD. In PEL, most cells express latent genes, with 1-5% of cells exhibiting spontaneous reactivation and lytic gene expression(7).

The mechanisms responsible for lytic reactivation in natural infection are unclear, however chemical inducers of lytic reactivation such as phorbol esters and histone deacetylase inhibitors have been utilized to study the lytic program of gene expression and more recently cell lines harboring latent virus and doxycycline inducible RTA provide a more specific model for the study of reactivation (8, 9). Aside from being the major switch for reactivation of KSHV from latency, RTA has intrinsic ubiquitin ligase activity and has been shown to interact with cellular ubiquitination machinery, ubiquitin ligases and deubiquitinases(10–12). RTA has also been designated a SUMO targeting ubiquitin ligase, specifically targeting SUMO2/3 modified proteins for proteasomal degradation(13, 14). RTA targets IRF7 for proteasomal degradation as a mechanism to abrogate the interferon α/β response to viral infection(11). In addition, RTA is reported to degrade a number of known RTA repressors such as ID2, MyD88, Hey1, LANA and NF-κB (p65)(15–20). This E3 ubiquitin ligase activity has been mapped to a Cys/His- rich domain between amino acids 118 and 207 of RTA(11). RTA was also shown to recruit and stabilize a cellular ubiquitin ligase, RAUL, via recruitment of the deubiquitinating enzyme HAUSP(10). We have previously reported RTA induced degradation of vFLIP through the cellular E3 ligase Itch (12, 21). We hypothesized that identification of additional substrates targeted for ubiquitination by RTA would provide insight into novel mechanisms of regulation of the latent-lytic transition. To this end, we carried out a comparative proteomics analysis in both RTA -transfected 293T cells and TREX BCBL-1 RTA SILAC labeled cells. We identified 66 ubiquitination sites in 40 proteins shared between both data sets, representing proteins with RTA -induced ubiquitination alterations in BCBL-1 cells. We evaluated RTA dependent modulation of protein abundance for a subset of the proteins in the context of a CURE (course-based undergraduate research experience). Undergraduate research experience is an important factor in persistence in STEM education and careers for students from diverse backgrounds (22). (22)(22) HLA-C, CDK1, MCM7, and SUMO2/3 were among the proteins identified as targets of RTA induced ubiquitination that displayed decreased protein abundance. Our dataset was enriched with proteins that are also known to be SUMOylated, with more than a third of the proteins in the matched dataset also reported to be SUMOylated. Inhibition of SUMOylation resulted in increased production of infectious virus, supporting a role for RTA induced degradation of SUMO in the transition to lytic replication. The ubiquitin ligase domains in RTA and RAUL were partially required for degradation of some of the substrates, suggesting that additional cellular factors are at play. We observed an RTA dependent decrease in the abundance of proteins associated with MHC class I antigen presentation and provide evidence supporting a novel mechanism of immune evasion associated with decreased TAP dependent peptide transport and intracellular HLA levels. This RTA dependent TAP2 inhibition represents a novel mechanism of attack on antigen presentation by KSHV early in lytic reactivation. In addition to identifying a novel mechanism of immune evasion, our study has yielded novel targets of RTA downmodulation that will enhance our understanding of the latent to lytic transition and provided 62 undergraduate students with authentic research experience.

## Methods

### Cell Lines, Plasmids, Transfection, and Antibodies

HEK 293T, iSLK, and BAC16 iSLK were cultured in DMEM medium supplemented with 10% FBS and TREx BCBL-1, iBC-3, and BC-3 cells were maintained in RPMI supplemented with 20% FBS. Cells were grown at 37°C with 5% CO2 under antibiotic selection where indicated (200ug/ml hygromycin). The following constructs were used in this study: FLAG tagged ORFs in pcDNA3.1+/C-(K)DYK were purchased from Genscript: CDK1 (NM_001786.5), MCM7 (NM_005916.5), VDAC1(NM_003374.3). HLAC (NM_002117.5) was obtained from Genecopoeia in a Gateway PLUS Shuttle clone and cloned into pcDNA-DEST40 (ThermoScientific). Flag tagged SUMO 2/3, RAUL (WT and C1051A) and RTA (WT and H145L) were provided by Diane and Gary Hayward. Cells were transfected using PEI (Polyethylenimine, Linear, MW 25000, Transfection Grade, Polysciences) at a ratio of 3ul PEL/1ug plasmid DNA. The following antibodies were used in this study: ß-actin (sc-69879), V5 (sc-271944) from Santa Cruz Biotech, FLAG M2 from Sigma Aldrich, FITC labeled HLA-ABC Antibody (11-9983-42 Invitrogen), PE Mouse Anti-Human CD184 (Clone 12G5, BD), Cdc2 p34 Sc-54 (Santa Cruz), HLA Class I ABC 15240-1-AP (Proteintech), MCM7 Sc-9966 (Santa Cruz), SUMO 2/3 AB374 (Abcam), RTA antibody was provided by Gary Hayward. Cells were transfected at 60-70% confluence using 1µg/ml polyethyleneimine (PEI) linear, MW∼25,000 at a ratio of 1µg plasmid DNA:3µl PEI. After 15 min of incubation at room temperature the mixture was added to the cells.

### Generation of iBC-3 stable cell line

Stable cells lines were generated using transduction with a lentiviral vector. Doxycycline inducible RTA (a gift from Prashant Desai, JHU) was selectable by hygromycin (200µg/ml). Lentiviral particles were generated by transfection of lentiviral packaging plasmids (PSPAX2 and PMD2.G) with transgene plasmid with PEI transfe193 ction reagent using protocol as recommended by manufacturer. Incubation lasted 18 hours before replacing media with SFM DMEM for 48 hours. Supernatant was collected and added dropwise to BC-3 cells, incubating for 24 hours before adding antibiotic for selection. Cells were evaluated for doxycycline induced RTA expression by western blot.

### Ubiquitin-modified proteome analysis

Proteomics experiments were conducted in collaboration with the Smoler Proteomics Center at the Technion-Israel Institute of Technology. Experiments were carried out using doxycycline inducible TREx BCBL-1 RTA cells and 293T cells transfected with RTA or pcDNA control vector. Cells were maintained as described above. Prior to ubiquitin remnant motif K-e-Gly-Gly (KGG) purification, cells were acclimated in SILAC media (lysine and arginine deficient media supplemented with stable isotope containing lysine and arginine (“heavy” media contained ^13^C6 ^15^N4 Arg and ^13^C6^15^N2 Lys)) for 7 days and evaluated for efficient SILAC labeling (23). RTA expression was induced in TREx BCBL-1 RTA cells via addition of doxycycline (1mg/ml) to cells for 24 and/or 48h as indicated. 293T cells were transfected with RTA or empty vector pcDNA 3.1 and harvested following 48h incubation. Cells were treated with 10mM MG132 6 h prior to harvesting. Cell pellets were washed with PBS and flash frozen in liquid nitrogen. Cells were lysed in 8M urea 400mM ammonium bicarbonate buffer pH 8, sonicated and cleared of cell debris by centrifugation. 2mg protein was processed for peptide purification by reducing with DTT (30min at 60°C, 3mM final concentration), cysteines were alkylated with 10mM idoacetamide, followed by trypsin digest (0.1mg/5ug protein) overnight. Following desalting by reverse phase chromatography on SepPac disposable cartridges (Waters), peptides containing the ubiquitin remnant motif – K-e-Gly-Gly were enriched via incubation with anti-K-e-Gly-Gly antibody-bound agarose beads. Peptides were eluted with 0.2% TFA. Eluted peptides were desalted using C18 stage tips and dried prior to analysis via LC-MS/MS.

The peptides were resolved by reverse-phase chromatography on 0.075 X 180-mm fused silica capillaries (J&W) packed with Reprosil reversed phase material (Dr Maisch GmbH, Germany). The peptides were eluted with linear 180 minutes gradient of 5 to 28% 15 minutes gradient of 28 to 95% and 25 minutes at 95% acetonitrile with 0.1% formic acid in water at flow rates of 0.15 μl/min. Mass spectrometry was performed by Q Executive HFX mass spectrometer (Thermo) in a positive mode using repetitively full MS scan followed by collision induces dissociation (HCD) of the 30 most dominant ions selected from the first MS scan.

MS raw data was processed using MaxQuant software followed by identification of proteins using the Andromeda peptide search engine to search against the HHV8 and human UniProt databases (Uniprot-proteome 27.7.18 (73101 entries), Uniprot-proteome HHV8) (24, 25). Data merging and statistical analysis was done using the Perseus(26). Functional enrichment analysis was performed using DAVID (https://david.ncifcrf.gov/) and STRING (https://string-db.org/). Final data set was curated by matching the data from the TREx BCBL-1 RTA dataset with the data from the 293T cell dataset to identify the RTA dependent ubiquitin-modified proteome. For time course proteomics experiments cells were cultured in the absence of MG132, lysates were separated on a 12% polyacrylamide gel and peptides extracted from Coomassie stained gel slices using in-gel digest and processed for LC-MS/MS without KGG purification. Experiments were performed in triplicate with a reverse SILAC labeling control, replicates with the highest Pearson’s correlation coefficient were used for the final data analysis.

### SDS-PAGE and Western Blot Analysis

Cells lysates were prepared by direct addition of 2X Laemmli buffer. Lysates were separated using a 4%-20% ExpressPlus PAGE Gel (Genescript) and MOPS running buffer. Proteins were transferred to a PVDF membrane using Trans-Blot Turbo using the Bio-Rad defined program for 1.5 mm gels. Membranes were blocked in 5% non-fat dry milk in PBS for 1 h followed by incubation with primary antibody overnight at 4°C and secondary antibody for 1 h at room temperature. Proteins were visualized by addition of ECL substrate and detection of chemiluminescence using a Licor C-DiGit or Azure 300.

### Quantitative real-time PCR

Cells were harvested and total RNA was isolated using GeneJet RNA purification kit. Poly A containing RNA was reverse transcribed to cDNA using a RevertAid First Strand cDNA Synthesis Kit per manufacturers protocol. The following primers were used to amplify the indicated genes and data was analyzed by the 2^^-^ΔΔ*Ct* method. Data obtained from >3 biological replicates was analyzed for significance by one-way ANOVA with Dunnett’s multiple comparisons test unless otherwise indicated in the figure legend.

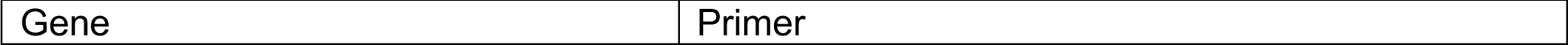
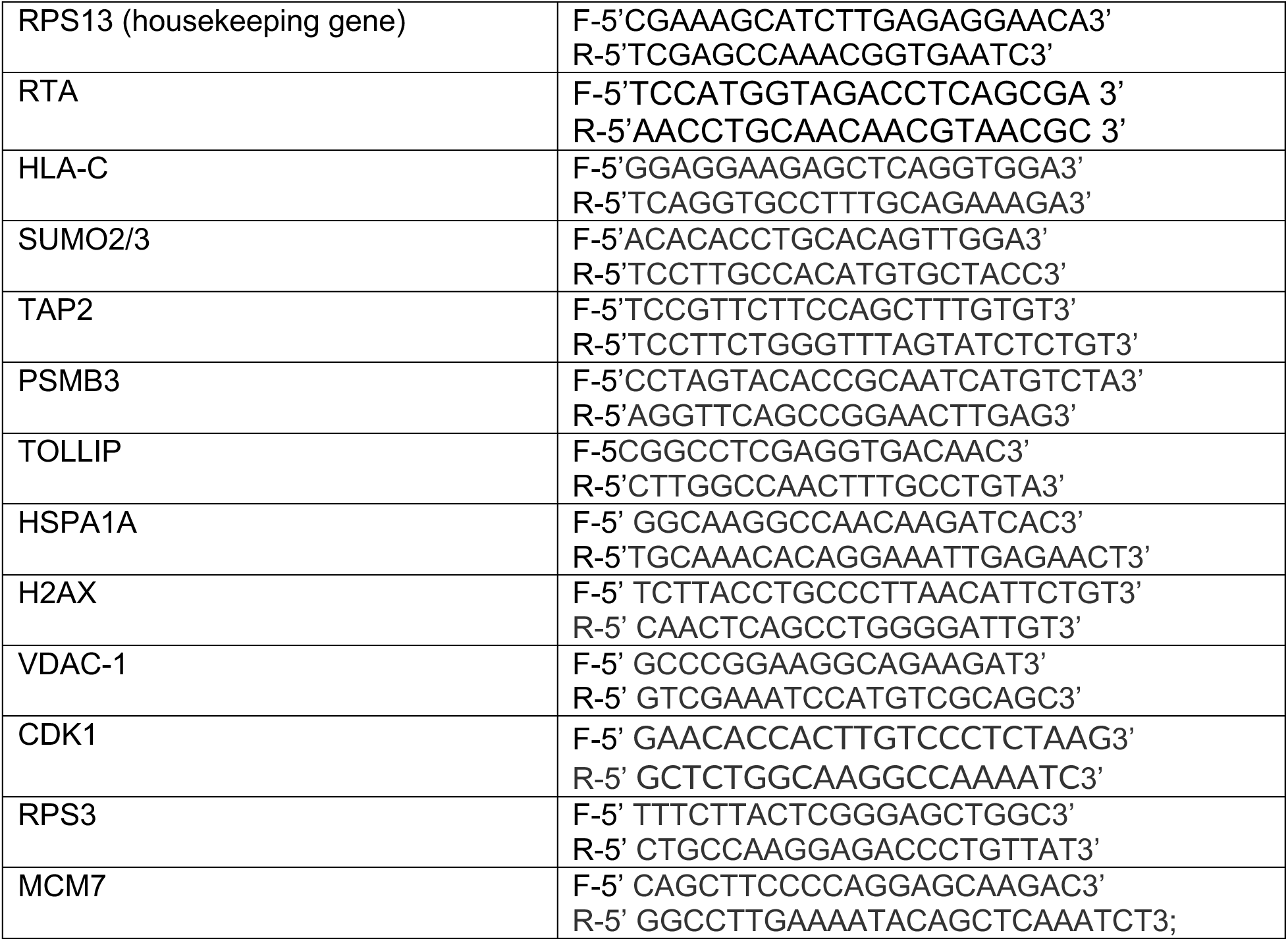

### TAP Peptide Transport Assay

iSLK cells were treated with 1ug/ml doxycycline at 90% confluence to induce RTA expression. 48 hrs. post doxycycline induction, cells were harvested, partially permeabilized and incubated with a peptide pool. Peptide pool contained the following cysteine AF 488 modified peptides at a concentration of 10nM: NST-F (RRYQNST[C]L), C4-F (RRY[C]KSTEL), and R9 (EPGY[C]NSTD).

Cells were incubated with 10mM ATP or ADP for 30 minutes at either 37°C or on ice. Cells were washed 3x with PBS before being fixed with 2% paraformaldehyde and analyzed using a Guava EasyCyte Flow cytometer.

### Surface and Intracellular Flow Cytometry

Cells were dissociated with 0.25% trypsin (Life Technologies) and resuspended in FACS wash buffer. Cells were washed twice, permeabilized (for intracellular staining only) and then incubated or directly incubated (surface staining only) with FITC labeled MHC ABC antibody (Invitrogen) or PE labled CXCR4 (BD Biosciences) at 4°C for 30 minutes. Cells were washed 3 times with FACS buffer at 4°C and analyzed using a Guava EasyCyte Flow cytometer.

### SeAP Assay

Vero rKSHV.294 and 293T MSR tet-OFF cells were kindly provided by David Lukac (27). Day one: Vero rKSHV.294 were plated in a six-well plate and treated with 2-D08, DMSO, or NaB where indicated. Day two: 293T MSR tet-OFF cells were plated in a six-well plate. Day three: media from 293T MSR tet-OFF cells was replaced with virus containing media from the rKSHV.294 cells. Day five: 25 ml of media was harvested and heat inactivated at 65°C for 30 minutes. SeAP activity was quantified using the Great EscAPe SeAP assay (Clontech). Fluorescence was read on a Filtermax F5 plate reader (Molecular Devices).

### Data availability

Raw data has been uploaded to the PRIDE database. Project Name: Analysis of the ubiquitin-modified proteome identifies novel host determinants of Kaposi’s sarcoma herpesvirus lytic reactivation Project accession: PXD046571.

## RESULTS

### Time-course of lytic induction

A proteomics experiment was first conducted in TREx BCBL-1 RTA cells to assess lytic gene expression over a 24h time-course. Peptides of viral origin displayed an increase in peptides of lytic origin and a sustained presence of other proteins such as LANA and vIL6 (Fig. 1a-b). In TREx BCBL-1 RTA cells, analysis of peptides of cellular origin revealed a decrease in the detection of proteins previously reported to be degraded following lytic induction (Fig. 1c). HLA-A/ HLA-B or MHC class I is known to be downregulated via the action of viral encoded ubiquitin ligases K3 and K5 (28–30). We also observed a decrease in BST2/Tetherin, also reported to be downregulated by K5 via endocytosis and subsequent downregulation in the endosome (31).

**Figure 1.**
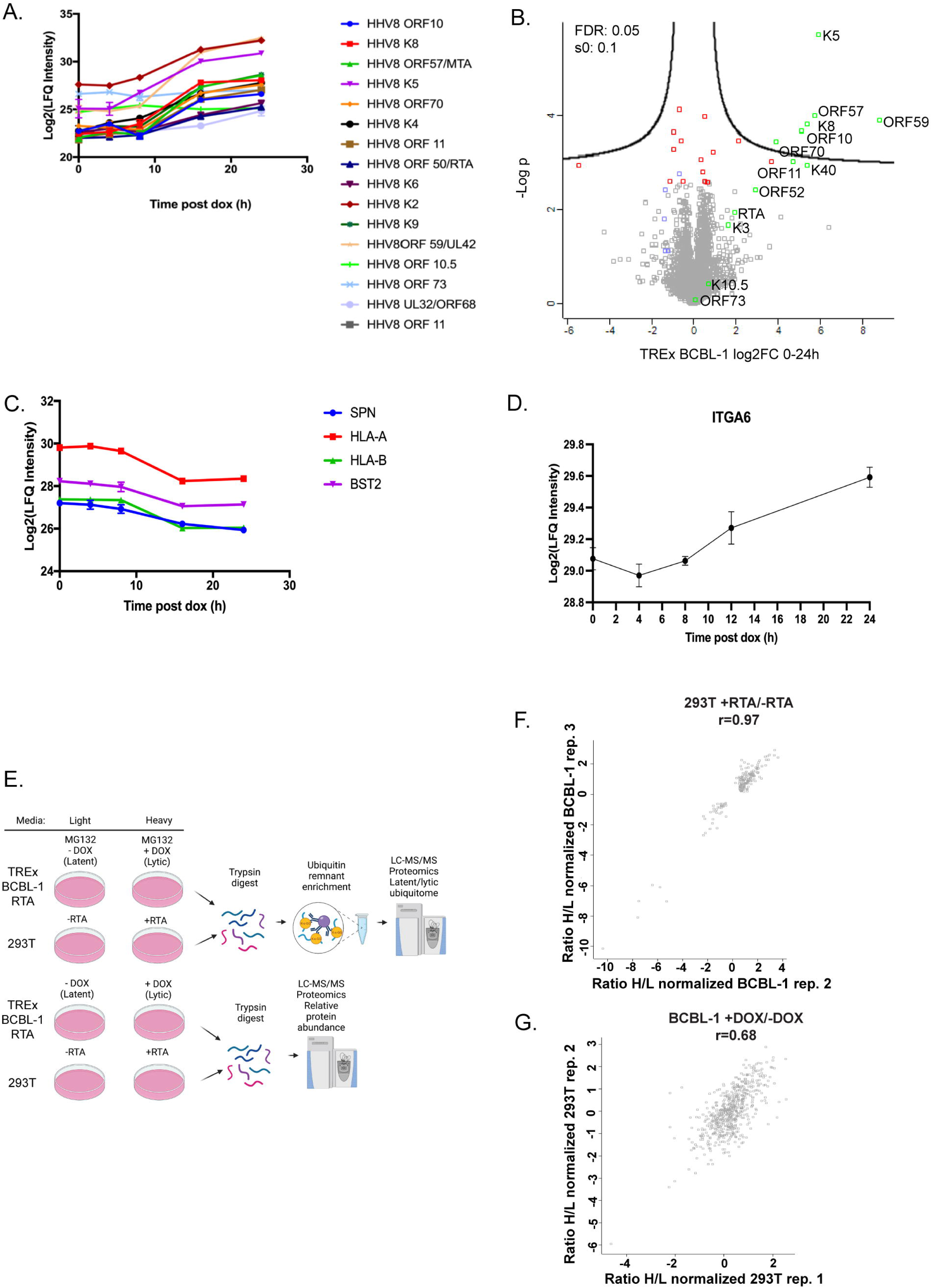
Time-course of lytic induction. (A-B) Doxycycline induced lytic reactivation of TREx BCBL-1 RTA cells resulted in (A) an increase in peptides derived from lytic proteins. (B) Volcano plot illustrating viral proteins in green and cellular proteins of interest noted in red and HLA is indicated in blue. (C) Peptides of cellular origin previously reported to be downregulated following lytic induction. (D) Integrin alpha 6 (ITGA6) displayed a modest increase. TREx BCBL-1 RTA cells were treated with doxycycline (1ug/ml) for 0, 4, 8, 16, and 24h. 293T cells were transfected with RTA using PEI. 1.8x107 cells were harvested in an 8M Urea buffer. 6ug of protein was reduced, alkylated, trypsin digested, cleaned on C18 stage tips and processed for MS. Data was analyzed using Maxquant, Perseus, and Graphpad Prism. (E) Schematic describing experimental design for comparative proteomics analysis. Experiments were carried out in both TREx BCBL-1 RTA and RTA transfected 293T cells. Cells were grown in SILAC media containing heavy or light lysine and arginine, supplemented with dialyzed FBS. Cells were treated with MG132 6h prior to harvest. 25ug of protein was saved for proteomic analyses following preparation of tryptic peptides. Tryptic peptides were enriched for ubiquitin remnants with anti-K-ε-GG antibody, followed by processing for mass spectrometry. Data was analyzed using Maxquant and Perseus. Created with Biorender.com. (F-G) Pearson correlation coefficients for biological replicates used in downstream analysis and experimentation. (F) RTA transfected 293T cells and (G) TREx BCBL- 1 RTA.

SPN/leukosialin was also significantly downregulated but has not been described previously (Fig. 1c). We also observed a modest but significant (q=0.012285714) increase in Integrin alpha-6 (Fig. 1d). This was interesting due to the involvement of integrins in cytomegalovirus entry and induction of angiogenesis (32, 33).

### Characterization of the RTA dependent KSHV ubiquitin-modified proteome

Given that RTA is the master lytic switch and has been shown to not only have intrinsic ubiquitin ligase and SUMO targeting ubiquitin ligase activity but is also reported to interact with cellular components of the ubiquitin proteasome system, we reasoned that identification of host proteins that displayed RTA dependent differential ubiquitination would increase our understanding of the latent to lytic transition. KSHV encodes two ubiquitin ligases (K3 and K5) that are expressed during lytic replication; however, our goal was to identify RTA specific targets. To identify targets of RTA induced differential ubiquitination, we designed our workflow in two disparate cell lines, doxycycline inducible TREx BCBL-1 RTA cells, which contain the entire viral genome, and RTA transfected 293T cells, which contain RTA only, with the rationale that the matched datasets would yield RTA specific targets. Cells were adapted to SILAC media as follows: light media contained “light” lysine and arginine and heavy media contained “heavy” lysine (13C6;15N2) and arginine (13C6;15N4) amino acids based on the protocol of Ong and Mann, 2007(23).

Following confirmation of stable labeling, cells were either treated with doxycycline to induce RTA expression or transfected with RTA or vector control. Experiments were conducted in each cell line in triplicate; twice where RTA was induced or transfected in “heavy” media labeled cells, and once where RTA was induced or transfected in “light” media labeled cells. Cells were either harvested for proteomics to evaluate lytic induction and protein abundance or treated with MG132 and incubated with anti-K-e-Gly-Gly antibody-bound agarose to isolate peptides containing ubiquitin remnants. Ubiquitinated peptides were enriched with anti-K-ε-GG antibody, followed by processing for mass spectrometry on a Q-Exactive HF-X Hybrid Quadrupole-Orbitrap Mass Spectrometer. For proteomics, to evaluate protein abundance, cells were harvested at the indicated time points, and lysates processed for MS. Data was analyzed using MaxQuant and Perseus. Experiments were performed in triplicate and the experiments with the highest Pearson correlation coefficient (293T RTA r=0.97 and TRExBCBL-1 RTA r=0.68) were used for further analysis (Fig.1e-g). (28–30)(31)(32, 33)

### Ubiquitin-modified proteome analysis in RTA transfected and naturally infected cells

Analysis of anti-K-ε-GG enriched ubiquitinated peptides resulted in the identification of 193 differentially ubiquitinated sites in 178 proteins in cells transfected with RTA, and 272 sites in 248 proteins in doxycycline treated TREx BCBL-1 RTA cells (Fig.2a). Sixty-six of these sites in 40 proteins were shared between both data sets, representing proteins with RTA-induced ubiquitination alterations in BCBL-1 cells; 29 displayed increased ubiquitination and 11 displayed decreased ubiquitination (Fig. 2b). While we observed greater variability within the TREx BCBL- 1 RTA experiment, we did observe an overall similar trend in the behavior of many ubiquitination sites in both cell lines (Fig. 2 d-e).

**Figure 2.**
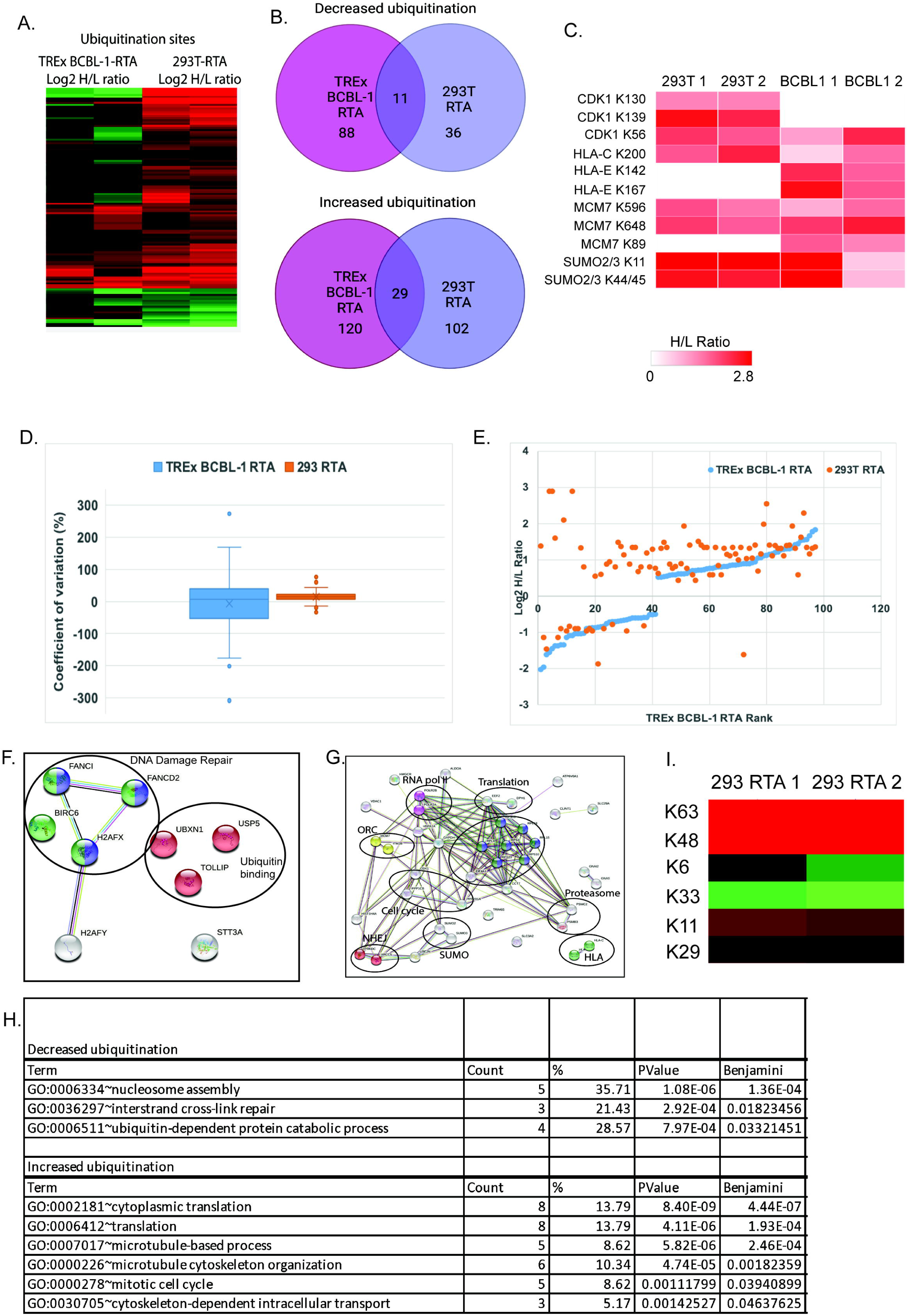
Analysis of the ubiquitin-modified proteome in RTA transfected and KSHV infected cells. (A) Heat map displaying shared ubiquitination sites between RTA transfected 293T and KSHV infected TREx BCBL-1 cells. The heat map was made from the matched matrix which contained both datasets and 207 peptides representing 98 unique proteins. Data was filtered to exclude peptides with a H/L ratio of -0.5 < x < 0.5. x < 0.5. Duplicate experiments are displayed for each cell type. Red=H/L ratio >0.5, Green=H/L ratio <0.5. (B) Venn diagram illustrating the number of proteins that displayed increased (top) and decreased (bottom) ubiquitination in each dataset and shared between datasets following filtering in Perseus. (C) Heat map illustrating shared ubiquitination sites that displayed increased ubiquitination in RTA-expressing cells. Heatmap generated by clustergrammer. (D) Percent coefficient of variation within each dataset. (E) Shared ubiquitination site scatterplot rank vs log2 H/L ratio demonstrating behavior of each modification in each dataset. Log2 H/L ratio >0.5, <-0.5 were considered significant increase or decrease respectively. Blue=TREx BCBL-1 RTA, Orange=293T RTA. (F-G) STRING analysis of proteins that displayed at least a 0.5-fold (F) decrease or (G) increase (Log2H/L ratio) in ubiquitination in the matched (BCBL-1 and 293T) dataset. (H) DAVID analysis of matched dataset highlighting significant GO terms. (I) K63 and K48 linked polyubiquitin chains were increased in RTA expressing 293T cells ((Log2H/L ratio = 0.57 and 1.49, respectively).

STRING analysis of proteins displaying an RTA dependent decrease in ubiquitination revealed two functional groups of proteins (Fig. 2f). One cluster is involved in DNA damage repair and the other is associated with ubiquitin binding. Analysis of proteins with increased ubiquitination in RTA expressing cells relative to control indicated functional clusters of proteins involved in diverse cellular processes including antigen presentation, DNA replication, DNA damage repair, cell cycle, SUMOylation, translation, transcription, and the proteasome (Fig. 2g) (34). Gene-annotation enrichment analysis using DAVID resulted in clustering into functional groups by gene ontology (GO term) (35, 36). The groups with the lowest Benjamini values are displayed in Fig. 2h. We also observed an increase in K48 and K63-linked polyubiquitin chains in RTA transfected 293T cells as well as a significant increase in ubiquitination of SUMO 2/3 in the matched dataset (Fig. 2i and Figure 8a). These observations are consistent with the reported functions of RTA as a ubiquitin ligase, manipulator of the ubiquitin proteasome system and SUMO targeting ubiquitin ligase (10–13).

### Evaluation of the abundance of selected cellular proteins in the presence of RTA

To validate the results of our proteomics experiment, we selected proteins from the matched dataset based on student generated hypotheses to evaluate protein stability in the presence of RTA expression. Flag tagged ORFs were expressed in 293T cells and evaluated for stability in the presence of RTA or pcDNA vector control. The following proteins that displayed increased ubiquitination in the ubiquitin-modified proteome screen, exhibited decreased abundance in the presence of RTA: HLA-C, CDK1, MCM7, VDAC1, HSPA1A, PSMB3, RPS3, and SUMO2/3 (Fig. 3a). H2AX displayed decreased ubiquitination in the proteomics and was stable in the presence of RTA, suggesting that impact of RTA on protein abundance was specific. Interestingly, TOLLIP displayed an RTA dependent decrease in ubiquitination in the ubiquitome and was drastically decreased in abundance in the presence of RTA, highlighting the broad functions of ubiquitination (Fig. 3a, see discussion). A decreased abundance of selected proteins was observed in doxycucline induced TREx BCBL-1 cells and in BC-3 cells induced for 24 or 48hr with TPA/NaB (Fig. 3b-c). We selected HLA-C, CDK1, SUMO2/3, and MCM7 for further analysis due to their reproducible RTA dependent decrease in protein abundance in PEL cells. To confirm that the observed decreased protein abundance was due to a post-transcriptional mechanism, we analyzed gene expression in both RTA transfected 293T and doxycycline induced BCBL-1 cells. We did not observe a significant decrease in gene expression in the presence of RTA suggesting a post-transcriptional mechanism at play (Fig. 4a-b). The loss of these proteins was dose-dependent and occurred rapidly as observed by cycloheximide chase (Fig. 5a-b).

**Figure 3.**
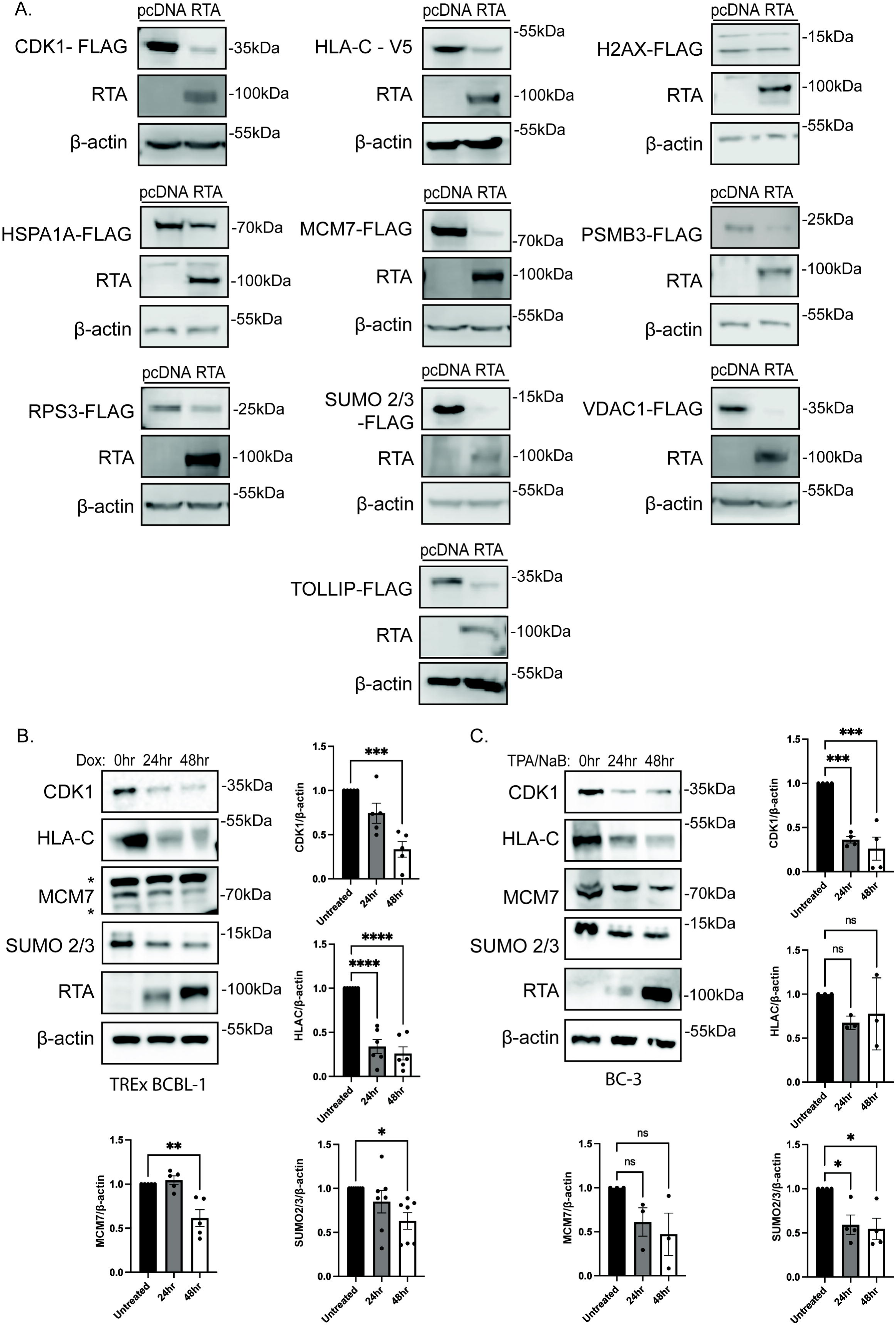
RTA expression reduces the abundance of selected proteins. (A) 293T cells were transfected with the indicated expression vectors in the absence or presence of RTA. Lysates were analyzed by immunoblot against the indicated epitope tags, RTA, or β-actin as a loading control. TREx BCBL-1 cells (B) and BC-3 cells (C) were treated with 1*μ*g/mL doxycycline or 50ng/mL TPA and 1mM sodium butyrate respectively. Lysates were collected at the indicated time points and analyzed by immunoblot for endogenous protein levels, RTA, and β-actin as a loading control. Representative immunoblot images are shown (left), * indicates non-specific bands. Quantification of abundance for indicated proteins (right/below) was performed using ImageJ software. Graphs represent the mean relative abundance of each protein compared to β- actin (n>3) ± SEM. Significance was calculated using one-way ANOVA with Dunnett’s multiple comparisons test. **** ≤0.0001, *** ≤0.001, ** ≤ 0.01, * <0.05

**Figure 4.**
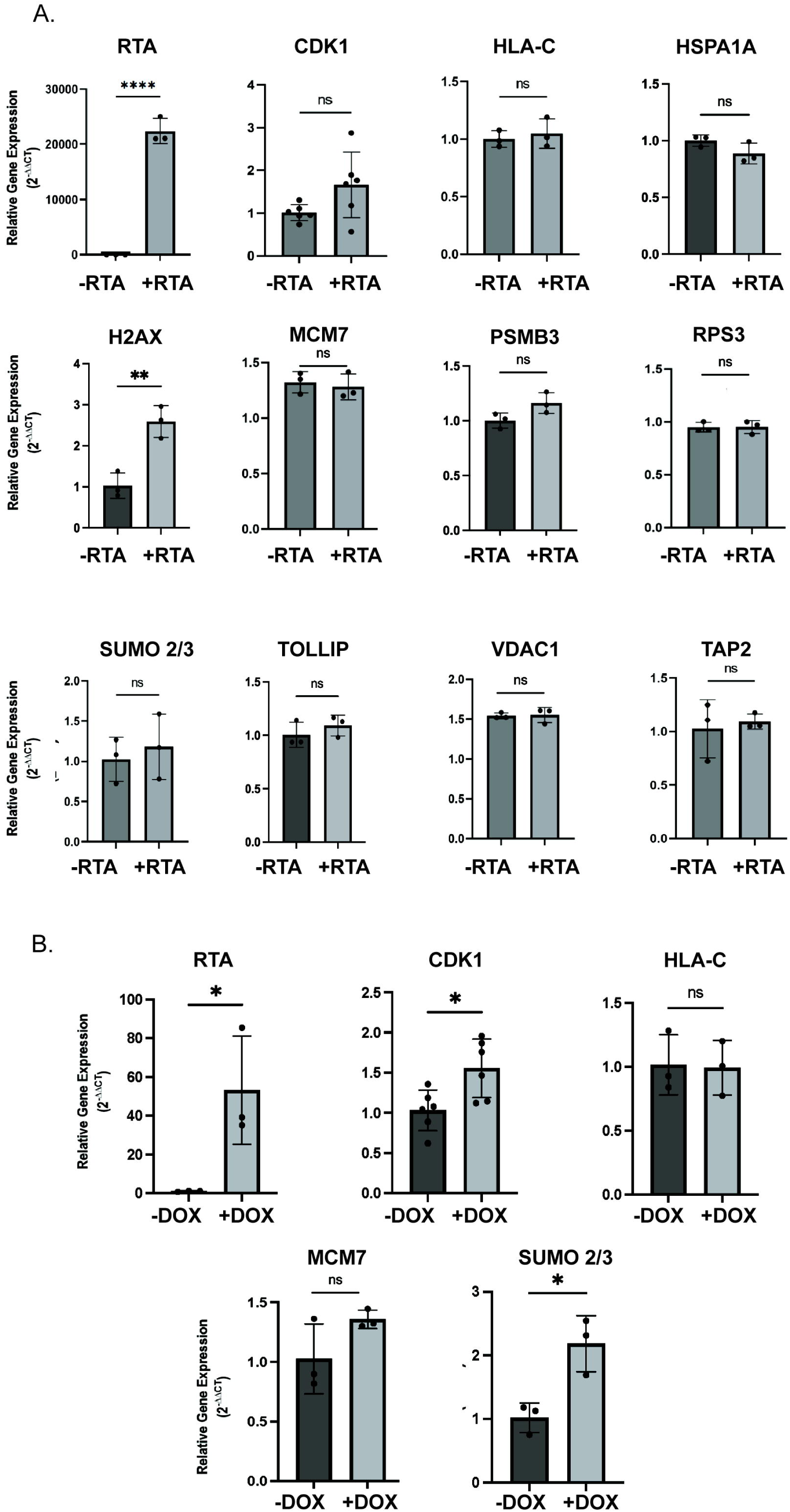
RTA does not reduce the expression of selected genes. (A) Transcript levels of target genes examined by qPCR in 293T cells transfected with or without RTA. (B) Transcript levels of target genes examined by qPCR in TREx BCBL-1 cells treated with 1ug/mL doxycycline for 24h. Relative gene expression was calculated using the 2^-ΔΔCT^ method relative to RPS13 housekeeping gene (n>3) ± SEM. Significance was calculated using one-way ANOVA with Dunnett’s multiple comparisons test. **** ≤0.0001, *** ≤0.001, ** ≤ 0.01, * <0.05

**Figure 5.**
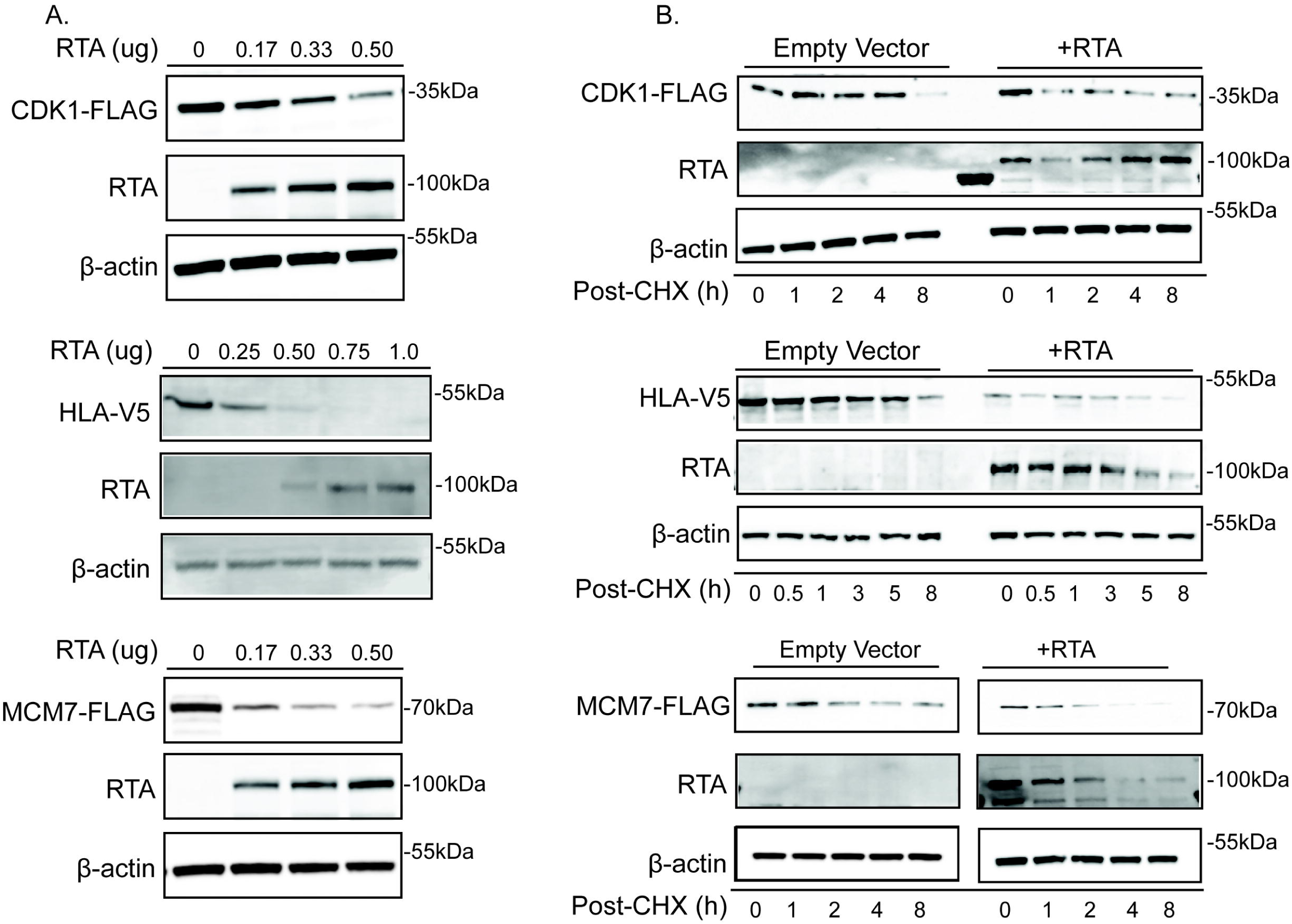
RTA rapidly reduces the abundance of selected proteins in a dose-dependent manner. (A) RTA induced degradation of selected targets is dose dependent. Increasing amounts of RTA were co-transfected with 1ug of indicated target gene in 293T cells. Lysates were analyzed by SDS-PAGE followed by immunoblot against FLAG or V5 epitope tag, RTA, or β-actin as a loading control. (B) Cycloheximide (CHX) chase illustrating RTA dependent reduction of target protein half-life. 293T cells were transfected with 1ug indicated tagged-protein and either empty vector or RTA. 24hr post-transfection, cycloheximide was added and cells were harvested at the indicated timepoints post CHX addition. Lysates were analyzed as described above.

### Characterizing the mechanism of RTA induced decrease of selected cellular proteins

RTA has ubiquitin ligase activity and interacts with cellular ubiquitin ligases. To further explore the mechanism responsible for the observed RTA dependent decrease in protein stability, we utilized ubiquitin ligase domain mutants of RTA, RAUL (UBE3C), and ITCH. RTA ubiquitin ligase activity has been mapped to an N-terminal RING-like domain (11). Substitution of leucine for histidine at position 145 has been shown to abolish ubiquitin ligase activity and stabilize substrates of RTA (11, 20, 37, 38). Mutations in this region have also been shown to significantly impede lytic replication. To determine whether RTA is destabilizing CDK1, HLA-C, MCM7, and/or SUMO2/3, 293T cells were transfected with epitope- tagged substrate and wild-type or mutant H145L RTA and protein abundance was observed. CDK1 was not stabilized in the presence of RTA H145L, however HLA-C, MCM7, and SUMO2/3 displayed varying levels of stability. HLA-C displayed the most evident stabilization, suggesting that RTA ubiquitin ligase activity is required for degradation (Fig. 6a). RTA is known to bind and stabilize the cellular ubiquitin ligase RAUL resulting in the degradation of RAUL substrates IRF-3 and IRF-7 (10). To evaluate whether RAUL ubiquitin ligase activity is required for degradation of CDK1, HLA-C, MCM7, and/or SUMO2/3, 293T cells were transfected with empty vector or RTA and wild-type or mutant RAUL C1051A and the indicated substrate. Wild-type and mutant RAUL expression had a stabilizing effect on MCM7 only (Fig. 6b). This result is counterintuitive as over-expression of wild-type RAUL should increase the degradation of the substrate. We discuss these findings further in the discussion. The Itch ubiquitin ligase, responsible for RTA induced degradation of vFLIP, was not responsible for the destabilization of any of the substrates evaluated (data not shown) (12). (11)(11, 20, 37, 38)(10)

**Figure 6.**
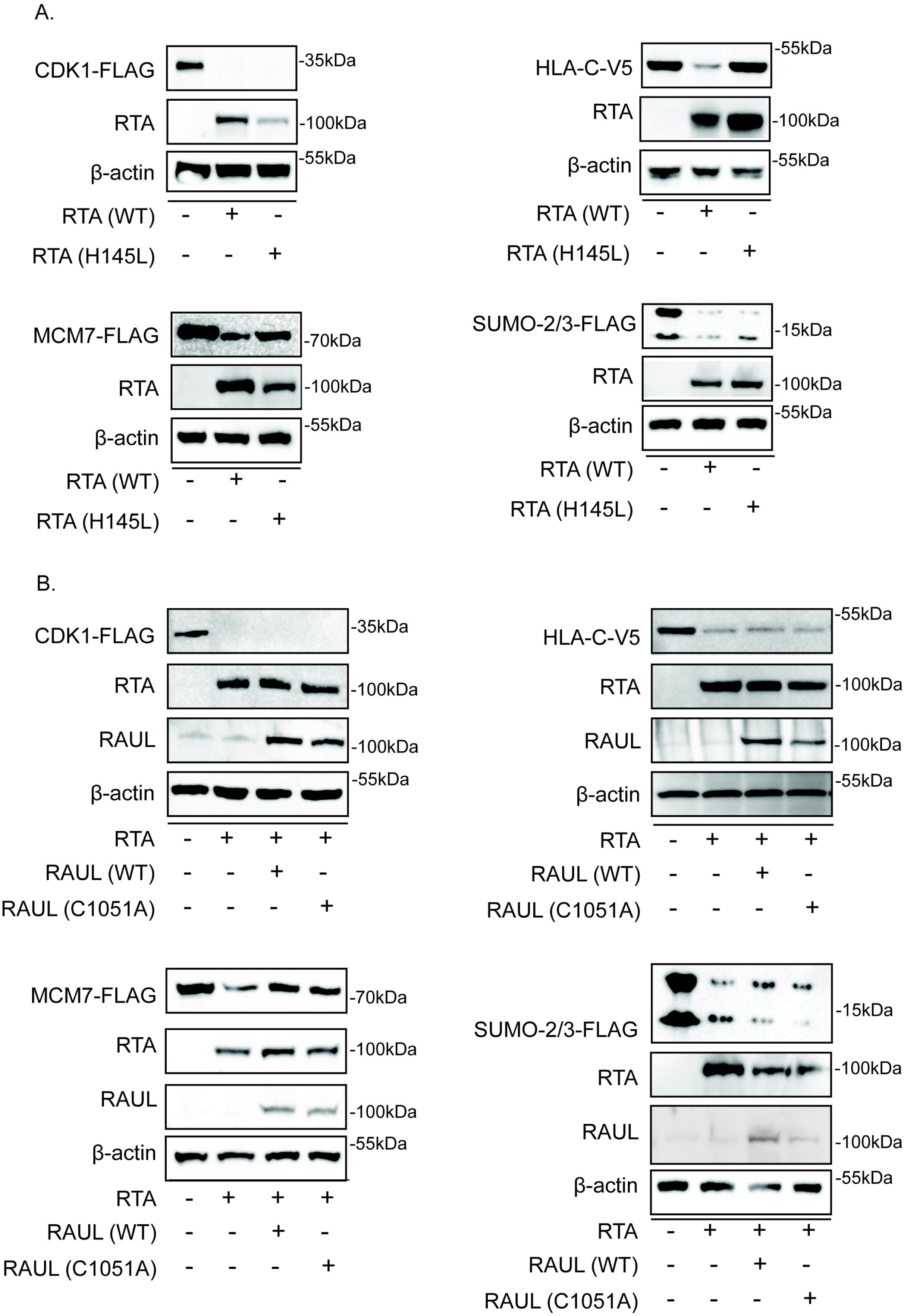
RTA targets cellular proteins for degradation through diverse mechanisms. (A) The RTA ubiquitin ligase domain is partially required for the degradation of some substrates of RTA. 293T cells were transfected with the indicated expression vector plus either wild-type or mutant (H145L) RTA or empty vector pcDNA control. Lysates were analyzed for substrate abundance via western blot. Results displayed are representative of at least three independent experiments. (B) The RAUL ubiquitin ligase domain is partially required for the RTA induced degradation of MCM7. 293T cells were transfected with the indicated substrate plus either wild-type or mutant (C1051A) RAUL or empty vector pcDNA control. Lysates were analyzed for substrate abundance via western blot. Results displayed are representative of at least three independent experiments.

### RTA targets SUMO2/3 for ubiquitination and degradation

RTA is a well characterized SUMO targeting ubiquitin ligase (STUbL) (13), therefore it was not surprising when SUMO 2/3 was identified as highly ubiquitinated at K11 and K45 (range 1.3-2.9 H/L ratio) in RTA expressing 293T and doxycycline induced TREx BCBL-1 RTA cells (Fig 7a). Further analysis of the ubiquitin-modified proteome dataset suggests that the RTA induced, or lytic, ubiquitin-modified proteome is enriched for proteins that are SUMOylated. According to PhosphoSitePlus, the human proteome contains approximately 12,434 proteins that are ubiquitinated, of those proteins, 2,434 or 19.5% are also reported to be SUMOylated, however within the RTA dependent ubiquitin-modified proteome 30.7% (TREx BCBL-1), 38.3% (293T), and 34.4% (matched matrix) of the proteins are also SUMOylated (Fig 3b). In fact, 61.5% of the proteins from the matched matrix that primarily localize to the nucleus are reported to be SUMOylated (Fig 7b). This observation suggests that the STUbL activity of RTA may be partly responsible for the observed increase in ubiquitinated proteins, specifically in the nuclear compartment. RTA is known to have intrinsic ubiquitin ligase activity as well as interact with cellular ubiquitin ligases and these, in addition to indirect mechanisms, may account for the remaining ubiquitination observed.

**Figure 7.**
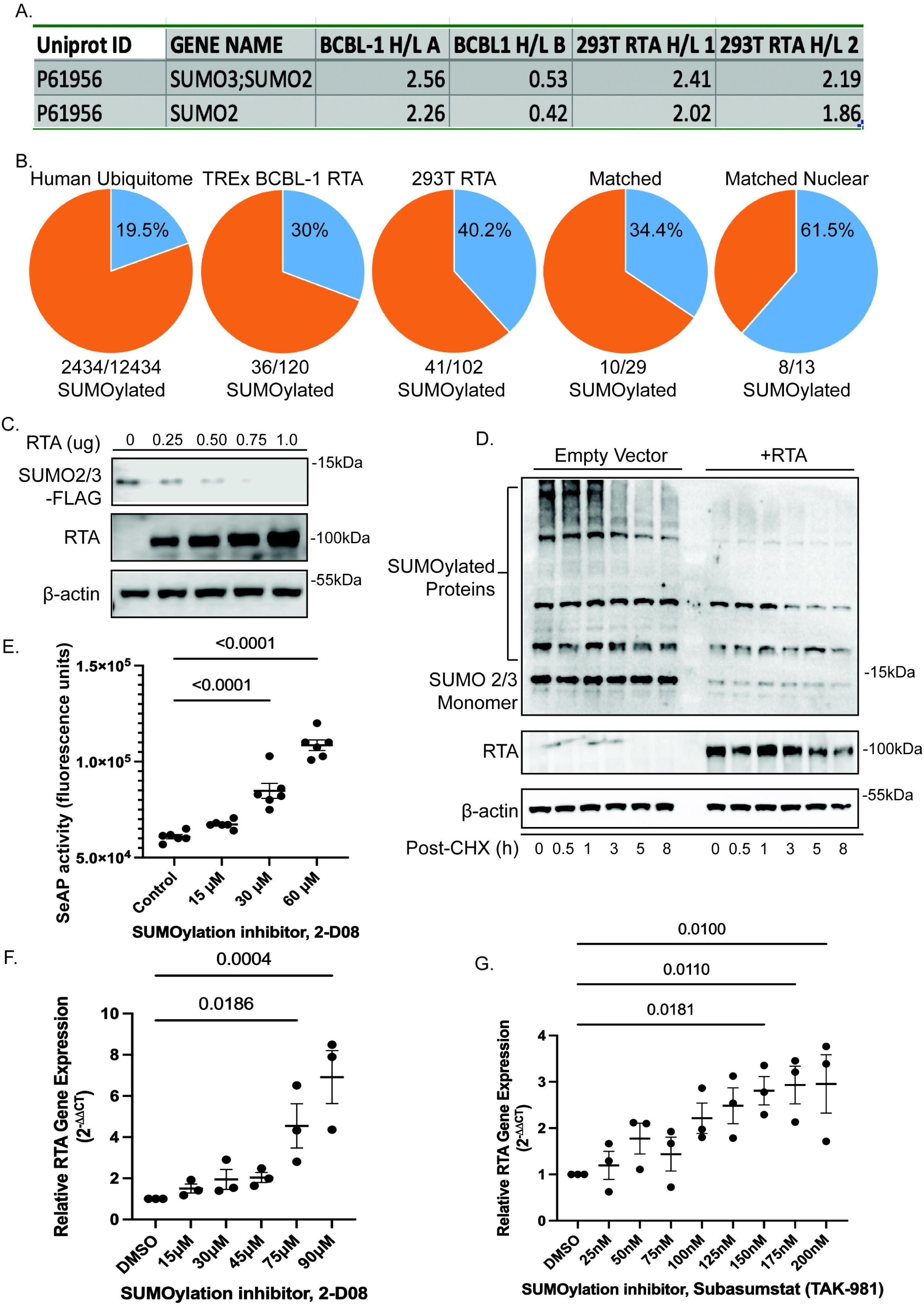
RTA targets SUMO 2/3 for degradation. (A) SUMO2/3 displayed increased ubiquitination in both doxycycline induced TREx BCBL-1 cells and RTA transfected 293T cells (Log2 H/L ratio). (B) The KSHV latent/lytic ubiquitin-modified proteome is enriched for proteins that are also SUMOylated. Proteins were evaluated for evidence of SUMOylation via PhosphoSitePlus. (C) RTA induced degradation of SUMO2/3 is dose dependent. Increasing amounts of RTA were co- transfected with 1ug of SUMO2/3 plasmid in 293T cells. Lysates were analyzed by SDS-PAGE followed by immunoblot against FLAG tag, RTA, or β-actin as a loading control. (D) Cycloheximide (CHX) chase illustrating RTA dependent reduction of SUMO2/3 half-life. 293T cells were transfected with 1ug SUMO2/3 expression vector and either empty vector or RTA. 24hr post- transfection, cycloheximide was added and cells were harvested at the indicated timepoints post CHX addition. Lysates were analyzed as described above. (E-G) Inhibition of SUMOylation results in lytic reactivation. (E) Vero rKSHV.294 cells that contain a recombinant KSHV genome expressing a SeAP reporter gene were treated with increasing concentrations of SUMOylation inhibitor 2-D08. Following 48h incubation, virus containing media was collected and added to 293 MSR Tet-OFF cells. Media was assayed for SeAP activity following 72h incubation. (F) iBC-3 cells were treated with increasing concentrations of SUMOylation inhibitor 2-D08. Following 48h incubation, cells were harvested, and RTA expression was analyzed by qPCR. (G) iBC-3 cells were treated with increasing concentrations of SUMOylation inhibitor Subasumstat (TAK-981). Following 48h incubation, cells were harvested, and RTA expression was analyzed by qPCR. Relative gene expression was calculated using the 2^-ΔΔCT^ method relative to RPS13 housekeeping gene (n=3 biological replicates) ± SEM. Significance was calculated using one-way ANOVA with Dunnett’s multiple comparisons test with p values displayed. Similar results were observed in parental BC-3 cells.

**Figure 8.**
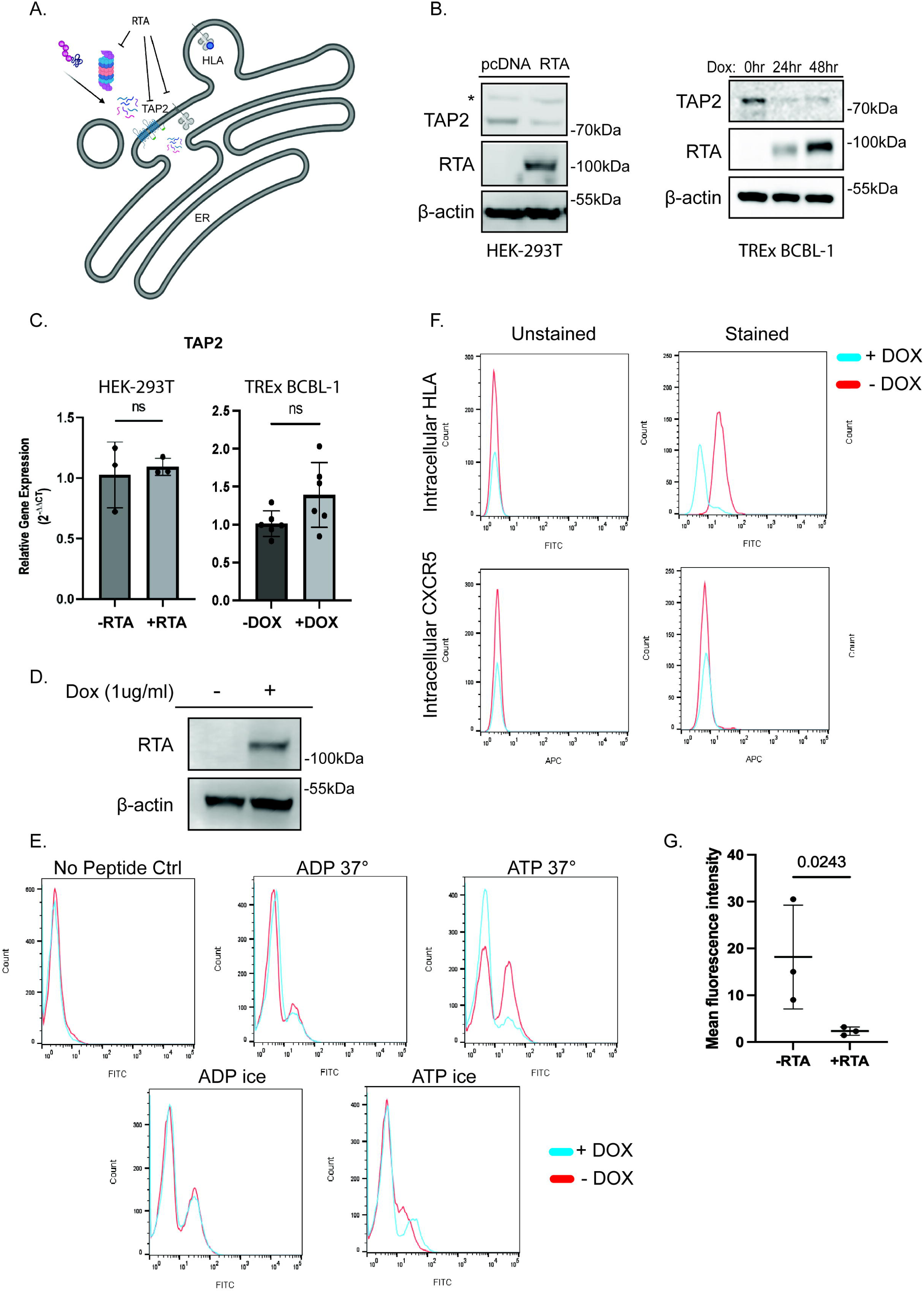
RTA targets antigen presentation by reducing peptide transport into the ER. (A) Proposed model in which RTA targets ER associated HLA by inhibiting TAP2. Created with Biorender.com. (B) RTA expression reduces the abundance of TAP2. 293T cells transduced with RTA (left) and BCBL1 cells treated with doxycycline to induce RTA (right) were harvested and lysates were analyzed by SDS-PAGE followed by immunoblot for TAP2, RTA, and β-actin. * Indicates non-specific bands. (C) Transcript levels of TAP2 examined by qPCR in 293T cells transfected with or without RTA (left) and in TREx BCBL-1 cells treated with 1ug/mL doxycycline for 24h (right). (D) RTA induction in iSLK cells. iSLK cells were treated ± 1ug/mL doxycycline for 24hr and harvested. Lysates were analyzed by SDS-PAGE followed by immunoblot for RTA and β-actin as a loading control. (E) Peptide transport assay. iSLK cells were treated ± doxycycline (1ug/mL) to induce RTA expression. 48hrs after treatment, cells were collected, semi- permeabilized and incubated with FITC labeled peptide pools under the indicated conditions. Cells were washed and FITC was quantified using a Guava EasyCyte flow cytometer. (F) RTA expression results in decreased intracellular HLA staining. iSLK cells were treated ± doxycycline (1ug/mL) to induce RTA expression. Cells were fixed, permeabilized and stained for HLA A,B,C or CXCR5 as a control. Left histograms = unstained cells, right = stained cells. Cells were analyzed as stated above. (G) Mean fluorescence intensity was analyzed for 3 replicate experiments using a ratio paired t test. P value = 0.0243

The RTA dependent degradation of SUMO 2/3 is dose dependent (Fig 7c) and occurs rapidly (Fig 7d). SUMOylation appears to play a role in maintaining latency, as inhibition of SUMOylation resulted in increased production of infectious virus as quantified by secreted alkaline phosphatase assay (Fig. 7e). We utilized Vero rKSHV.294 cells that contain a reporter gene that expresses secreted placental alkaline phosphatase (SeAP) from a tetracycline responsive promoter. SeAP is secreted into the media only after successful infection of 293 MSR Tet OFF cells. Vero rKSHV.294 cells were treated with increasing concentrations of SUMOylation inhibitor, 2-D08. After 48h incubation, virus containing media was collected and added to 293 MSR Tet OFF cells. Media was assayed for SeAP activity following 72h incubation. We observed a significant dose dependent increase (p<0.0001) in SeAP activity at noncytotoxic concentrations of 30µM and higher. Similarly, inhibition of SUMOylation with 2-DO8 or the highly selective SUMOylation inhibitor Subasumstat (TAK-981) resulted in a dose-dependent increase in RTA induction in iBC- 3 cells (Fig. 7f-g). These data suggest that RTA targets SUMO 2/3 for degradation and that SUMOylation is important for maintaining latency in Vero rKSHV.294 and BC-3 cells.

### RTA targets antigen presentation by targeting peptide import into the ER

Analysis of the ubiquitin-modified proteome yielded two proteins that are associated with antigen presentation: HLA (A, B and C) and TAP2. These proteins play a role in antigen processing and presentation (Fig. 8a). We initially observed a decrease in HLA A and B abundance in the time course of doxycycline induced RTA expression in BCBL-1cells (Fig. 1c). While KSHV K3 and K5 are known to downregulate MHC class I via endocytosis (39), ubiquitination was observed in RTA transfected 293T cells in addition to doxycycline treated TREx BCBL-1 cells, suggesting that there may be an additional RTA dependent mechanism for HLA downregulation (Fig. 2 c). TAP2 is an ABC transporter that, in complex with TAP1, participates in the transport of proteasome generated peptides from the cytosol to the endoplasmic reticulum (ER) for loading onto nascent MHC class I (HLA) molecules. The TAP1-TAP2 peptide transport complex is targeted by multiple viruses, serving as an effective mechanism of immune evasion. TAP2 is reported to interact with Epstein Barr virus BLNF2a, herpes simplex virus US12/ICP47, and adenovirus E3-19K (40–43). Through mass spectrometry, we observed TAP2 ubiquitination at K356 which is located within the peptide binding loop. We observed an RTA dependent decrease in HLA abundance following 24h RTA expression (Fig. 3a-c). This degradation was dose dependent with RTA expression reducing HLA half-life to 2h post cycloheximide addition compared to 6h in empty vector transfected cells (Fig. 5a-b). Similarly, we observed a decrease in endogenous TAP2 abundance in both 293T and TREx BCBL-1 cells (Fig. 8b).

Based on these observations, we hypothesized that peptide transport was being inhibited through depletion of TAP2 and this was resulting in decreased HLA abundance. To evaluate this hypothesis, we employed a peptide transport assay based on the method described by Fishbach et al. (44). These experiments were carried out in iSLK cells lacking BAC16, allowing us to separate the function of RTA from the function of KSHV encoded genes like K3 and K5. This assay utilized a pool of FITC-labeled peptides. Doxycycline inducible iSLK RTA cells lacking BAC16 were subjected to mild permeabilization and incubated with FITC labeled peptide pools under the indicated conditions. This allowed for plasma membrane permeabilization, however the ER membrane remained intact. Following 30 min incubation with labeled peptide pools, cells were washed to remove any peptide not actively transported into the ER, and peptide translocation was quantified by flow cytometric detection of FITC. We observed active translocation of peptides into the ER in iSLK cells incubated at 37°C in the presence of ATP, however cells with doxycycline induced RTA expression (Fig. 8d) displayed decreased peptide translocation under the same conditions, suggesting that RTA is interfering with TAP dependent transport of peptides into the ER (Fig. 8e). Proteins that are not properly folded in the ER are targeted for degradation by ERAD. Specifically, HLA molecules that fail to bind *β*-2-microglobulin or peptide are retained, retrotranslocated, ubiquitinated and degraded (45). Mass spectrometry analysis of HLA ubiquitination induced by *β*-2-microglobulin depletion identified ubiquitination at two lysines located in the ER lumen, K200 and K267(46). We observed RTA induced ubiquitination of HLA-B and C at K200 via mass spectrometry, suggesting that the ubiquitination is occurring due to the absence of peptide. If inhibition of peptide transport and subsequent degradation of HLA via ERAD were to occur, we would expect to see decreased intracellular staining of HLA. iSLK cells lacking BAC16 were treated with doxycycline or media control followed by staining for endogenous intracellular HLA A, B, and C or CXCR5 as a control. RTA expressing cells displayed significantly decreased (9-fold average decrease in mean fluorescence intensity) intracellular HLA staining compared to untreated controls. There was no impact on the unrelated receptor CXCR5 (Fig. 8f-g). Taken together, these data support our hypothesis that KSHV RTA targets antigen presentation through inhibition of peptide import into the ER resulting in a decrease in HLA biosynthesis/maturation.

## Discussion

RTA is the primary regulator of the latent to lytic transition in KSHV. RTA is not only the transcriptional transactivator responsible for the activation of lytic gene expression, but also has ubiquitin ligase activity and is known to both directly and indirectly target proteins of viral and cellular origin for ubiquitination. Many of the targets of RTA induced degradation represent repressors of lytic reactivation. Targets of RTA include SMC5/6, ID2, SUMO 2/3, vFLIP, IRF7, MyD88, Hey1, LANA and NFκB (p65) (10, 12, 13, 15, 17–19, 38, 47, 48) Using the rationale that RTA targets repressors of lytic reactivation we set out to identify novel regulators of the latent to lytic transition using a proteomics-based approach. We utilized SILAC labeling and ubiquitin remnant enrichment to identify proteins that display RTA dependent differential ubiquitination. We designed our experiment in two disparate cell lines; RTA transfected 293T and doxycycline induced BCBL-1, allowing us to merge our data sets to identify proteins that displayed RTA dependent differential ubiquitination and omit K3 and K5 dependent ubiquitination. We used datasets with the highest correlation coefficients for downstream analyses (RTA transfected 293T r=0.97, doxycycline induced TREx BCBL-1 r=0.67). We noted that the correlation coefficient was lower for doxycycline induced TREx BCBL-1 samples compared to RTA transfected 293T cells and hypothesize that this may be due to the complexity of the system. With doxycycline induced lytic reactivation, there are the added factors of doxycycline associated gene expression, lower RTA expression in BCBL-1 compared to transfected 293T, the effects of viral ubiquitin ligases K3 and K5, in addition lytic gene expression. We attempted to neutralize the effects of this variability by filtering out any proteins that did not display similar ubiquitination activity in replicates and matching the remaining proteins with the RTA transfected 293T dataset. We did observe some variability when validating the data in PEL cells and used this step to further refine our “hits”. Some proteins, like MCM7, were completely destabilized 100% of the time in transfected 293T cells, while the degradation was more variable in PEL cells. We illustrate this variability through quantification of all (≥3) biological replicates.

We observed an RTA dependent increase in K-48 and K-63 linked polyubiquitin chains. We identified 40 proteins that displayed RTA dependent differential ubiquitination. These proteins function in cellular processes including DNA damage repair, transcriptional regulation, DNA replication, apoptosis, and immune evasion. Through the development of a course-based undergraduate research experience (CURE), designed to provide authentic research experiences to undergraduate students, we selected proteins to validate and further characterize. Students selected proteins, generated hypotheses, and validated degradation of TAP2, HLA-C, CDK1, MCM7, VDAC1, HSPA1A, PSMB3, RPS3, and SUMO2/3. In a subsequent semester, students asked whether RTA is directly ubiquitinating these substrates and utilized a ubiquitin ligase domain mutant of RTA to evaluate protein stability. Participation in this course over a three-year period exposed 62 undergraduate students to the process of research and inspired a few of them to consider biomedical research as a career path.

In this study we present the following observations: 1. RTA expression results in cell specific reprogramming of the ubiquitome 2. RTA targets the degradation of SUMOylated proteins to promote lytic replication and 3. RTA expression decreases TAP2 protein levels abrogating peptide transport into the ER resulting in decreased intracellular HLA.

RTA is a well characterized STUbL, so it was not surprising when we observed SUMO 2/3 to be among the most highly ubiquitinated proteins identified in our RTA dependent ubiquitin-modified proteome(13). In fact, our ubiquitome was enriched for ubiquitinated proteins that are reported as SUMOylated via Phosphosite Plus, often on the same lysine. When we analyzed the SUMOylation status and cellular localization of the proteins that displayed an increase in ubiquitination, we found that 61.5% of the of the proteins that were also SUMOylated are localized to the nucleus. RTA is primarily localized to the nucleus. Taken together, this observation supports a model where nuclear localized RTA targets SUMOylated proteins to promote lytic reactivation. Recently the nuclear localization of RTA was reported to be regulated by linear ubiquitination on K516 by cellular ubiquitin ligase HOIP (49). It would be interesting to see whether mutation of this lysine impacts the stability of nuclear targets of RTA. To probe the functional role of SUMOylation in maintaining latency we utilized the infectivity assay developed by the Viera and Lukac labs(27, 50) When Vero rKSHV.294 cells, containing virus with the gene encoding secreted alkaline phosphatase, were treated with increasing doses of a small molecule inhibitor of SUMOylation, we observed a dose dependent increase in production of infectious virus. We saw a similar increase in lytic induction when BC-3 cells were treated with increasing concentrations of SUMOylation inhibitor 2-DO8. These data are consistent with reports that RTA STUbL activity is required for lytic reactivation and gene expression (13).

To explore the mechanism of RTA induced degradation of selected proteins, we asked whether the ubiquitin ligase domains of RTA, RAUL, and Itch are required for degradation. The RTA ubiquitin ligase domain appears to be required for degradation of most of the proteins evaluated, however we did not observe 100% recovery to control levels. When we evaluated the role of RAUL ubiquitin ligase activity, we only observed stabilization with MCM7. We observed stabilization of MCM7 in the presence of RTA and wild-type and dominant negative mutant RAUL. This finding was both surprising and counterintuitive, as one would predict degradation in the presence of wild-type RAUL. We have observed in previous work with the Cul5 ubiquitin ligase that overexpression of any one protein in this multiprotein ubiquitin ligase results in a dominant negative effect. We hypothesize that this is due to the disruption of the stoichiometry of the complex. Over expression of wild-type Cul5, Elongin B or Elongin C, resulted in a loss of ubiquitin ligase activity. We hypothesize that a similar effect is happening here, that RAUL and RTA work with an additional protein(s) to destabilize MCM7 and over expression of 2 components the complex results in formation of multiple incomplete complexes, resulting in a defect in ubiquitin ligase activity and enhanced protein stability. This observation leads to the hypothesis that RAUL may regulate MCM7 stability in the absence of RTA. Further experiments need to be carried out to test these hypotheses. (10, 12, 18, 19)

KSHV encoded ubiquitin ligases K3 and K5 play an important role in immune evasion via ubiquitination and subsequent endocytosis of HLA(39). We observed increased ubiquitination and degradation of HLA in both doxycycline induced TREx BCBL-1 RTA and RTA transfected 293T cells in addition to decreased levels of peptides of HLA origin in TREx BCBL-1 cells as early as 12hrs post lytic induction. Ubiquitination and subsequent degradation of HLA-C was observed in RTA transfected 293T cells, suggesting an additional, K3/K5 independent, mechanism of HLA targeted immune evasion. DNA viruses have evolved diverse mechanisms to target antigen presentation, frequently targeting the TAP peptide transporter. Herpesviruses in particular frequently target TAP function. Cytomegalovirus US6 targets TAP by inhibiting ATP binding(42, 51). HSV-1 ICP47 inhibits peptide transport by blocking the peptide-binding site and EBV interferes with both peptide and ATP binding (40, 43). We observed TAP2 ubiquitination at K356 which is located within the peptide binding loop(52). Expression of RTA in iSLK cells resulted in reduced peptide transport. The exact mechanism of inhibition of peptide transport remains unclear, but potentially include RTA induced degradation of TAP2 or ubiquitination of TAP2, resulting in decreased peptide binding.

RTA is primarily localized to the nucleus. This begs the question of how RTA is targeting cytoplasmic proteins. 13/29 of the proteins that displayed increased ubiquitination are nuclear, but 16 are localized to the cytoplasm. RTA is translated in the cytoplasm and imported into the nucleus and recently the mechanism for regulation of RTA localization was elucidated (49). It is possible that some activity occurs prior to nuclear import or that a subset of RTA localizes to the cytoplasm. This model is supported by our observation that the RTA has the largest impact on HLA stability 24h post transfection/doxycycline induction, with a decreased effect at 48h when RTA has presumably localized to the nucleus. Alternatively, RTA may be exerting its effects on cytoplasmic proteins indirectly, either through increasing expression of cellular ubiquitin ligases or destabilizing a deubiquitinase that is both nuclear and cytoplasmic. In the case of TAP2, this complex is localized to the ER membrane. The ER membrane is contiguous with the nuclear envelope, increasing the potential for interactions while the RTA is being imported into the nucleus. It is likely that aside from the SUMO targeting activity of RTA, one model will not explain how all proteins are targeted and this remains an area for future research.

Taken together, here we have identified 40 proteins that display RTA dependent differential ubiquitination. We validated 17.2% of the proteins that displayed increased ubiquitination for RTA dependent degradation. Our dataset was enriched with proteins that are also known to be SUMOylated. Inhibition of SUMOylation resulted in increased production of infectious virus, supporting a role for RTA induced degradation of SUMO and SUMOylated proteins in the transition from latency to lytic replication. We observed RTA dependent degradation of proteins associated with MHC class I antigen presentation and provide evidence supporting a novel mechanism of immune evasion associated with decreased TAP dependent peptide transport and increased intracellular HLA and proteasome subunit degradation. This report contributes to our understanding of KSHV biology by identifying new points of virus-host interaction and describing a novel mechanism of immune evasion by KSHV RTA during lytic replication that likely contributes to the success of the virus in establishing a lifelong infection. Future study of the RTA targets identified in this study have the potential to further our understanding of the latent to lytic transition and lead to the identification of new chemotherapeutics.

## Acknowledgements

We acknowledge the students from BIOL412 the KSHV Ubiquitome CURE for their help in conducting many of the experiments in this study.

## References

1. Dittmer DP, Damania B. 2013. Kaposi sarcoma associated herpesvirus pathogenesis (KSHV)--an update. Curr Opin Virol 3:238–44.

2. Dissinger NJ, Damania B. 2016. Recent advances in understanding Kaposi’s sarcoma-associated herpesvirus. F1000Res 5:740.

3. Bower M, Nelson M, Young AM, Thirlwell C, Newsom-Davis T, Mandalia S, Dhillon T, Holmes P, Gazzard BG, Stebbing J. 2005. Immune reconstitution inflammatory syndrome associated with Kaposi’s sarcoma. Journal of Clinical Oncology 23:5224–5228.

4. Uldrick TS, Wang V, Mahony DO, Aleman K, Wyvill KM, Marshall V, Steinberg SM, Pittaluga S, Maric I, Whitby D, Tosato G, Little RF, Yarchoan R. 2010. An Interleukin-6 Related Systemic Inflammatory Syndrome in Patients Co-Infected with Kaposi Sarcoma Associated Herpesvirus and HIV but without Multicentric Castleman Disease 1868:350– 358.

4. Polizzotto MN, Uldrick TS, Hu D, Yarchoan R. 2012. Clinical manifestations of Kaposi sarcoma herpesvirus lytic activation: Multicentric Castleman disease (KSHV-MCD) and the KSHV inflammatory cytokine syndrome. Front Microbiol 3:1–9.

5. Https://ntp.niehs.nih.gov/go/roc14 N (National TP. 2016. Report on Carcinogens, Fourteenth Edition. Research Triangle Park, NC: U.S.

6. Cesarman E, Damania B, Krown SE, Martin J, Bower M, Whitby D. 2019. Kaposi sarcoma. Nat Rev Dis Primers 5:9.

7. Nakamura H, Lu M, Gwack Y, Souvlis J, Zeichner SL, Jung JU. 2003. Global Changes in Kaposis Sarcoma-Associated Virus Gene Expression Patterns following Expression of a Tetracycline-Inducible Rta Transactivator. J Virol 77:4205 LP – 4220.

8. Myoung J, Ganem D. 2011. Generation of a doxycycline-inducible KSHV producer cell line of endothelial origin: Maintenance of tight latency with efficient reactivation upon induction. J Virol Methods 174:12–21.

9. Yu Y, Hayward GS. 2010. The ubiquitin E3 ligase RAUL negatively regulates type i interferon through ubiquitination of the transcription factors IRF7 and IRF3. Immunity 33:863–77.

10. Yu Y, Wang SE, Hayward GS. 2005. The KSHV immediate-early transcription factor RTA encodes ubiquitin E3 ligase activity that targets IRF7 for proteosome-mediated degradation. Immunity 22:59–70.

11. Chmura JC, Herold K, Ruffin A, Atuobi T, Fabiyi Y, Mitchell AE, Choi YB, Ehrlich ES. 2017. The Itch ubiquitin ligase is required for KSHV RTA induced vFLIP degradation. Virology 501.

12. Izumiya Y, Kobayashi K, Kim KY, Pochampalli M, Izumiya C, Shevchenko B, Wang DH, Huerta SB, Martinez A, Campbell M, Kung HJ. 2013. Kaposi’s Sarcoma-Associated Herpesvirus K-Rta Exhibits SUMO-Targeting Ubiquitin Ligase (STUbL) Like Activity and Is Essential for Viral Reactivation. PLoS Pathog 9.

13. Campbell M, Izumiya Y. 2012. Post-translational modifications of Kaposi’s sarcoma- associated herpesvirus regulatory proteins - SUMO and KSHV. Front Microbiol 3:1–13.

14. Cai Q-L, Knight JS, Verma SC, Zald P, Robertson ES. 2006. EC5S ubiquitin complex is recruited by KSHV latent antigen LANA for degradation of the VHL and p53 tumor suppressors. PLoS Pathog 2:e116.

15. Zhao Q, Liang D, Sun R, Jia B, Xia T, Xiao H, Lan K. 2015. Kaposis Sarcoma-Associated Herpesvirus-Encoded Replication and Transcription Activator Impairs Innate Immunity via Ubiquitin-Mediated Degradation of Myeloid Differentiation Factor 88. J Virol 89:415 LP – 427.

16. Yang Z, Yan Z, Wood C. 2008. Kaposis Sarcoma-Associated Herpesvirus Transactivator RTA Promotes Degradation of the Repressors To Regulate Viral Lytic Replication. J Virol 82:3590 LP – 3603.

17. Gould F, Harrison SM, Hewitt EW, Whitehouse A. 2009. Kaposis Sarcoma-Associated Herpesvirus RTA Promotes Degradation of the Hey1 Repressor Protein through the Ubiquitin Proteasome Pathway. J Virol 83:6727 LP – 6738.

18. Combs LR, Spires LM, Alonso JD, Papp B, Toth Z. 2022. KSHV RTA Induces Degradation of the Host Transcription Repressor ID2 To Promote the Viral Lytic Cycle. J Virol 96.

19. Combs LR, Combs J, McKenna R, Toth Z. 2023. Protein Degradation by Gammaherpesvirus RTAs: More Than Just Viral Transactivators. Viruses 10.3390/v15030730.

20. Ehrlich ES, Chmura JC, Smith JC, Kalu NN, Hayward GS. 2014. KSHV RTA abolishes NF B responsive gene expression during lytic reactivation by targeting vFLIP for degradation via the proteasome. PLoS One 9.

21. Bangera G, Brownell SE. 2014. Course-based undergraduate research experiences can make scientific research more inclusive. CBE Life Sci Educ 13.

22. Ong SE, Mann M. 2007. A practical recipe for stable isotope labeling by amino acids in cell culture (SILAC). Nat Protoc 1:2650–2660.

23. Cox J, Mann M. 2008. MaxQuant enables high peptide identification rates, individualized p.p.b.-range mass accuracies and proteome-wide protein quantification. Nat Biotechnol 26:1367–1372.

24. Cox J, Neuhauser N, Michalski A, Scheltema RA, Olsen J v., Mann M. 2011. Andromeda: A peptide search engine integrated into the MaxQuant environment. J Proteome Res 10:1794–1805.

25. Tyanova S, Temu T, Sinitcyn P, Carlson A, Hein MY, Geiger T, Mann M, Cox J. 2016. The Perseus computational platform for comprehensive analysis of (prote)omics data. Nat Methods 10.1038/nmeth.3901.

26. DeCotiis JL, Ortiz NC, Vega BA, Lukac DM. 2017. An easily transfectable cell line that produces an infectious reporter virus for routine and robust quantitation of Kaposi’s sarcoma-associated herpesvirus reactivation. J Virol Methods 247:99–106.

27. Coscoy L, Ganem D. 2002. Kaposi’s sarcoma-associated herpesvirus encodes two proteins that block cell surface display of MHC class I chains by enhancing their endocytosis. Proceedings of the National Academy of Sciences 97:8051–8056.

28. Parkinson MDJ, Piper SC, Bright NA, Evans JL, Boname JM, Bowers K, Lehner PJ, Luzio JP. 2015. A non-canonical ESCRT pathway, including histidine domain phosphotyrosine phosphatase (HD-PTP), is used for down-regulation of virally ubiquitinated MHC class I. Biochemical Journal 471:79–88.

29. Goto E, Yamanaka Y, Ishikawa A, Aoki-Kawasumi M, Mito-Yoshida M, Ohmura- Hoshino M, Matsuki Y, Kajikawa M, Hirano H, Ishido S. 2010. Contribution of Lysine 11-linked Ubiquitination to MIR2-mediated Major Histocompatibility Complex Class I Internalization. Journal of Biological Chemistry 285:35311–35319.

30. Pardieu C, Vigan R, Wilson SJ, Calvi A, Zang T, Bieniasz P, Kellam P, Towers GJ, Neil SJD. 2010. The RING-CH Ligase K5 Antagonizes Restriction of KSHV and HIV-1 Particle Release by Mediating Ubiquitin-Dependent Endosomal Degradation of Tetherin. PLoS Pathog 6:e1000843.

31. Campadelli-Fiume G, Collins-McMillen D, Gianni T, Yurochko AD. 2016. Integrins as Herpesvirus Receptors and Mediators of the Host Signalosome. Annu Rev Virol 3:215– 236.

32. Feire AL, Koss H, Compton T. 2004. Cellular integrins function as entry receptors for human cytomegalovirus via a highly conserved disintegrin-like domain.

33. Szklarczyk D, Gable AL, Lyon D, Junge A, Wyder S, Huerta-Cepas J, Simonovic M, Doncheva NT, Morris JH, Bork P, Jensen LJ, Mering C von. 2018. STRING v11: protein– protein association networks with increased coverage, supporting functional discovery in genome-wide experimental datasets. Nucleic Acids Res 47:D607–D613.

34. Huang DW, Sherman BT, Lempicki RA. 2008. Bioinformatics enrichment tools: paths toward the comprehensive functional analysis of large gene lists. Nucleic Acids Res 37:1– 13.

35. Huang J, Wang Y, Guo Y, Sun S. 2010. Down-regulated microRNA-152 induces aberrant DNA methylation in hepatitis B virus-related hepatocellular carcinoma by targeting DNA methyltransferase 1. Hepatology 52:60–70.

36. Gould F, Harrison SM, Hewitt EW, Whitehouse A. 2009. Kaposi’s sarcoma-associated herpesvirus RTA promotes degradation of the Hey1 repressor protein through the ubiquitin proteasome pathway. J Virol 83:6727–38.

37. Zhao Q, Liang D, Sun R, Jia B, Xia T, Xiao H, Lan K. 2015. Kaposi’s Sarcoma-Associated Herpesvirus-Encoded Replication and Transcription Activator Impairs Innate Immunity via Ubiquitin-Mediated Degradation of Myeloid Differentiation Factor 88. J Virol 89:415–427.

38. Ishido S, Wang C, Lee B-S, Cohen GB, Jung JU. 2000. Downregulation of Major Histocompatibility Complex Class I Molecules by Kaposis Sarcoma-Associated Herpesvirus K3 and K5 Proteins. J Virol 74:5300 LP – 5309.

39. Ressing Kremmer ME, Rickinson AB, Wiertz EJHJ, Garstka MA, Hislop AD, Horst ED, van Leeuwen D, Croft NP. 2009. Antigen Presentation Impairment of HLA Class I-Restricted Associated with Antigen Processing Results in Protein BNLF2a to the Transporter Specific Targeting of the EBV Lytic Phase 10.4049/jimmunol.0803218.

40. Bennett EM, Bennink JR, Yewdell JW, Brodsky FM. 1999. Cutting edge: adenovirus E19 has two mechanisms for affecting class I MHC expression. J Immunol 162.

41. Ahn K, Gruhler A. 1997. The ER-Luminal Domain of the HCMV Glycoprotein US6 Inhibits Peptide Translocation by TAPImmunity.

42. Ahn K, Meyer TH, Uebel S, Sempé P, Djaballah H, Yang Y, Peterson PA, Früh K, Tampé R. 1996. Molecular mechanism and species specificity of TAP inhibition by herpes simplex virus protein ICP47. EMBO Journal 15:3247–3255.

43. Fischbach H, Döring M, Nikles D, Lehnert E, Baldauf C, Kalinke U, Tampé R. 2015. ARTICLE Ultrasensitive quantification of TAP-dependent antigen compartmentalization in scarce primary immune cell subsets 10.1038/ncomms7199.

44. Hughes EA, Hammond C, Cresswell P, Amos DB. 1997. Misfolded major histocompatibility complex class I heavy chains are translocated into the cytoplasm and degraded by the proteasome.

45. Burr ML, van den Boomen DJH, Bye H, Antrobus R, Wiertz EJ, Lehner PJ. 2013. MHC class I molecules are preferentially ubiquitinated on endoplasmic reticulum luminal residues during HRD1 ubiquitin E3 ligase-mediated dislocation. Proc Natl Acad Sci U S A 110:14290–14295.

46. Lubyova B, Kellum MJ, Frisancho AJ, Pitha PM. 2004. Kaposi’s Sarcoma-associated Herpesvirus-encoded vIRF-3 Stimulates the Transcriptional Activity of Cellular IRF-3 and IRF-7. Journal of Biological Chemistry 279:7643–7654.

47. Han C, Zhang D, Gui C, Huang L, Chang S, Dong L, Bai L, Wu S, LanID K. 2022. KSHV RTA antagonizes SMC5/6 complex-induced viral chromatin compaction by hijacking the ubiquitin-proteasome system 10.1371/journal.ppat.1010744.

48. Luan Y, Long W, Dai L, Tao P, Deng Z, Xia Z. 2024. Linear ubiquitination regulates the KSHV replication and transcription activator protein to control infection. Nat Commun 15:5515.

49. Gantt S, Carlsson J, Ikoma M, Gachelet E, Gray M, Geballe AP, Corey L, Casper C, Lagunoff M, Vieira J. 2011. The HIV protease inhibitor nelfinavir inhibits Kaposi’s sarcoma-associated herpesvirus replication in vitro. Antimicrob Agents Chemother 55:2696–2703.

50. The EMBO Journal - 2001 - Hewitt - The human cytomegalovirus gene product US6 inhibits ATP binding by TAP.

51. Hewitt EW, Lehner PJ. The ABC-transporter signature motif is required for peptide translocation but not peptide binding by TAP.

